# Scaling the fitness effects of mutations with respect to differentially adapted *Arabidopsis thaliana* accessions under natural conditions

**DOI:** 10.1101/2023.10.12.561936

**Authors:** Frank W. Stearns, Juannan Zhou, Charles B. Fenster

## Abstract

Mutations are the ultimate source of genetic variation for natural selection to act upon. A major question in evolutionary biology is the extent to which new mutations can generate genetic variation under natural conditions to permit adaptive evolution over ecological time scales. Here we collected fitness data for chemically induced (ethylmethane sulfonate, EMS) mutant lines descended from two *Arabidopsis thaliana* ecotypes that show differential adaptation to the local environment of our common garden plot. Using a novel nonparametric Bayesian statistical approach, we found that both ecotypes accumulated substantial proportions of beneficial mutations. The poorly adapted ecotype showed higher variance in the fitness effect of mutations than the well-adapted ecotype. Furthermore, we predict that it takes less than 4000 generations for the fitness space of the two ecotypes to overlap through mutation accumulation, and that a single founder, through mutation accumulation, is able to achieve the species-wide genetic variation in less than 10,000 generations. Our results provide evidence for relatively rapid local adaptation of *Arabidopsis thaliana* in natural conditions through new mutations, as well as the utility of nonparametric Bayesian method for modeling the distribution of fitness effects for field-collected data.

## Introduction

Since mutations are the ultimate source of genetic variation, quantifying mutation parameters is fundamental to understanding evolutionary processes such as adaptation genetics (Lynch and Walsh 1998). Key mutation parameters are the distribution of mutation effects on fitness (DFE) and the magnitude of fitness effects of mutations (Eyre-Walker and Keightley 2007; Halligan and Keightley 2009). Because the vast majority of new mutation studies have been conducted on microorganisms under laboratory conditions (Eyre-Walker and Keightley 2007) we lack estimates of these parameters in macro-organisms under natural conditions where the effects of mutations can be assessed in the environment where natural selection occurs. Whether mutations have beneficial effects on fitness traits rarely or more frequently and the size of these effects scaled to environmental effects on the trait(s) will determine the role of new mutations in a population’s response to selection. This has implications for use in crop production (e.g., Chaudhary et al. 2019; Gaines et al. 2020) and whether populations can evolve in place in response to the stresses imposed by anthropogenic alteration of the environment, including climate change (Frankham et al. 2017).

One approach to studying mutations is spontaneous mutation accumulation. Mutation accumulation lines (MA lines) have been used since at least 1928 (Muller) to study the cumulative effects of mutations on traits, including fitness (Halligan and Keightley 2009). MA lines are lines that are derived from a common nearly homozygous founder, that are cultivated via intense inbreeding (full sib mating if separate sexes, or selfing, if hermaphrodites) and thus, genetically diverge through the accumulation of independent mutations (Lynch and Walsh 1998). Using the MA line approach, lab-based estimates of the contribution of mutations to heritable genetic variance (h^2^_m_, Lynch and Walsh 1998; Shaw et al. 2000) are typically 10-fold higher than field-based estimates (Rutter et al. 2010, 2018; Roles et al. 2016), with 500 and 5000 generations based on lab and field-based estimates, respectively, needed for mutation to generate a typical h^2^ of 50%. This reflects the much greater environmental variance in field versus lab settings.

While field-based estimates of h^2^_m_ are low, a number of studies indicate that mutations have the potential to contribute to population genetic variation in a meaningful way. First, field-based studies with *Arabidopsis thaliana* reveal that a significant proportion of new mutations are beneficial (Rutter et al. 2010, 2018; Roles et al. 2016; Weng et al. 2021) although how specific mutations affect fitness is dependent on the environmental context (Stearns and Fenster 2016a,b; Rutter et al. 2018; Weng et al. 2021). Second, the amount of genetic variation generated by mutations can, in a relevant number of generations, generate levels of genetic variation in *A. thaliana* seen within and between populations (Rutter et al. 2018; Weng et al. 2021). Third, there is a direct link between the likelihood of a site mutating and whether that site is polymorphic in natural populations of *A. thaliana* (Weng et al. 2019; Monroe et al. 2022), suggesting that such studies are biologically meaningful.

The above evidence suggest that mutations have the potential to contribute to the evolution of adaptations through their response to selection at microevolutionary time scales. However, the use of MA lines to represent new mutations and the subsequent analyses limit inference of mutational input to adaptive response. Even using the model system *A. thaliana*, with generation times of 3-4 per year, it is difficult to produce many generations of mutation accumulation. All field studies to date to assess the fitness effects of mutations in *A. thaliana* have been limited to 8-25 generations of mutation accumulations. Second, previous studies of mutation accumulation, whether in the field (above citations) or lab (Shaw et al. 2002; Rutter et al. 2010), infer the mutation effect on fitness by comparing the distribution of line fitness relative to the founder. Parametrizing mutation effects have required the DFE to be within a designated family of probability distributions (e.g. Gamma distribution) and impose prior distributions (e.g. Shaw et al. 2002; Eyre-Walker and Keightley 2009; Rutter et al. 2010). This practice relies on strong assumptions of the shape of the DFE, and therefore could potentially introduce bias. Finally, the contribution of mutation to population differentiation scaled to actual population genetic differentiation has been inferred (Rutter et al. 2018) and only rarely directly included in experimental designs (Weng et al. 2021). Consequently, we have limited understanding of the potential contribution of individual mutations to the patterns of genetic differentiation commonly observed in nature.

Here we use a novel nonparametric Bayesian hierarchical modeling approach to estimate the DFE and magnitude of mutation effects under field conditions for two ecotypes of *Arabidopsis thaliana* using chemically induced (ethylmethane sulfonate, EMS) mutation lines and four reference ecotypes with no associated mutant lines. Our approach allows us to avoid potential bias to determine the effect of starting fitness on frequency and size of beneficial mutations. We also calibrate the effect of mutations in fitness space by testing whether mutations alone can change the performance of one accession to be equivalent to another of higher performance. Finally, our use of EMS lines allows us to quantify the total effects of many more mutations relative to MA line approaches relying on new mutational events. Sampling many simultaneous mutations may allow us to better estimate the distribution of effects of mutations because we are able to make assumptions in our modeling approach by exploiting the Central Limit Theorem. Consequently, this approach enables us to estimate the mean and standard deviation of the DFE without making a priori assumptions about the shape of the DFE. Our Bayesian model also relies on several other innovative approaches, including accounting for covariance structures between sublines due to segregating mutations, as well as the usage of a logistic function to model survival probability as a function of genetic values, allowing us to simultaneously fit all fitness data that contains many zeros due to non-surviving seedlings.

## Methods

### Mutagenesis

Mutagenesis followed the methods from Stearns and Fenster (2016b). Briefly, seeds were collected from a single individual for each accession. Seeds were then treated with a 20 μM solution of ethylmethane sulfonate (EMS), an alkylating agent that induces G:C to A:T substitutions (Greene et al., 2003). The spectrum of mutations produced by EMS is similar to that of spontaneous mutations, although it does not include indels (Greene et al. 2003). Based on the proportion of the albino seed phenotype, we estimated 25 mutations per cell in coding regions per genome (Stearns and Fenster 2016b). This mutation rate resulted in an approximate six-fold greater number of nonsynonymous mutations in protein coding sequence (Weng et al. 2019) as quantified from direct sequencing of Columbia mutation accumulation lines representing 25 generations of spontaneous MA (Ossowski et al. 2010). We therefore expect our lines to carry the number of mutations roughly equivalent to 150 generations of mutation accumulation. Mutant and founder lines were established from single seeds with/without EMS treatment. The lines were raised in growth chamber conditions (Stearns and Fenster 2016b) and allowed to self-fertilize for two generations. They were then split into two sublines to account for maternal effects and allowed to self-fertilize for an additional generation to produce the seeds for plants used in this experiment (M4 generation) (Fig. 1).

**Figure 1.**
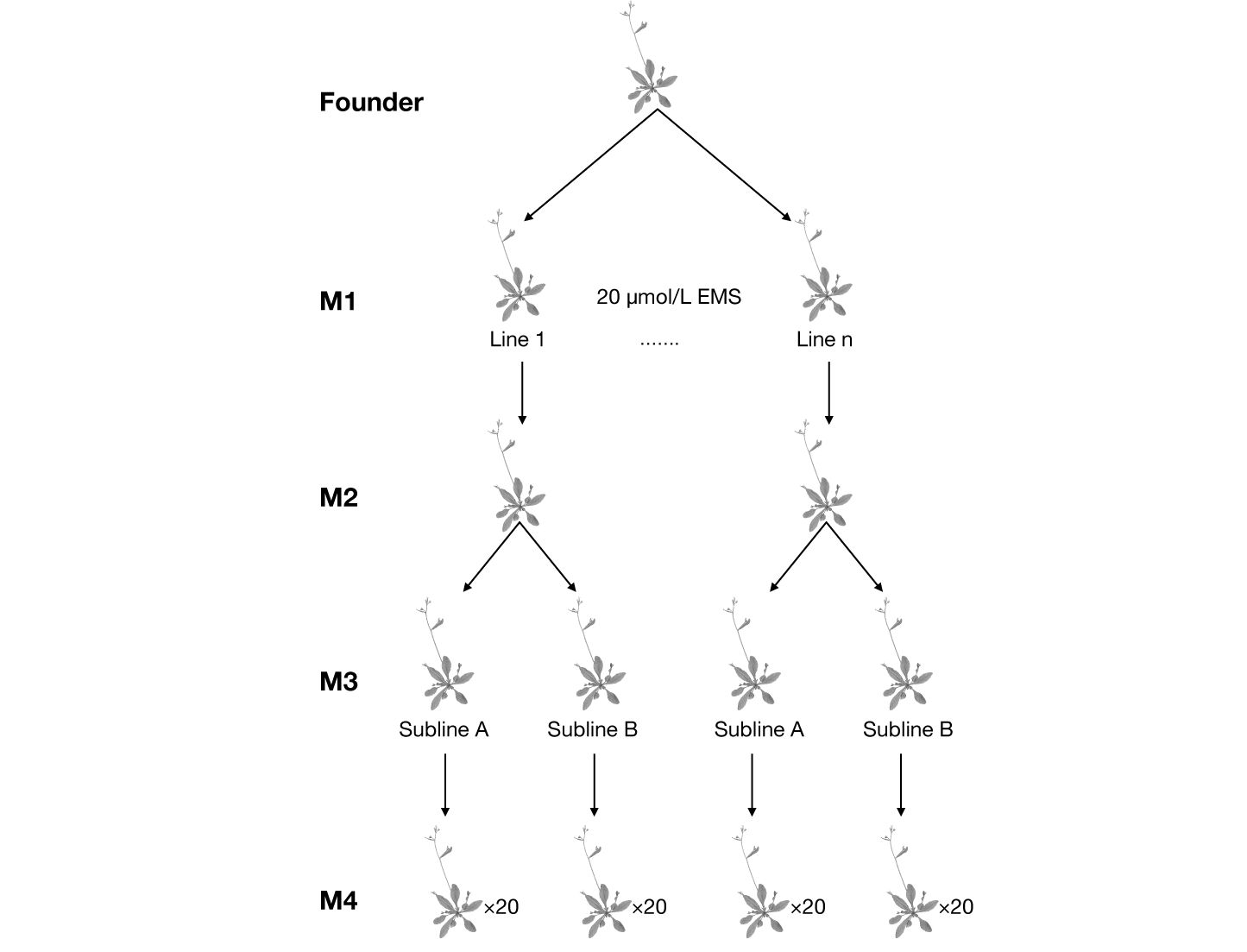
A schematic representation showing the process of generating mutation sublines from *Arabidopsis thaliana*. Seeds were collected from a single Columbia founder and treated with 20 μmol/L ethylmethane sulfonate (EMS) for 12 hrs. Twenty and 16 mutant lines were derived from this treatment for the Columbia and Cape Verde founders, respectively. Each line was then split into two sublines each (M3 generation), and seeds from these sublines (via selfing) were planted in the field (Beltsville Experimental Agricultural Station (UMD) in Beltsville MD (N 39.05378 W −76.95387).

### Fitness Assessment

All seedlings were planted at the Beltsville Agricultural Station (UMD) in Beltsville, MD on one day during the Fall of 2014. The timing of planting corresponds to the predominant winter annual life cycle that is exhibited by *A. thaliana* throughout its range (fall germination, overwinter as a vegetative rosette, spring flowering and senescence). Average low temperatures are several degrees below freezing for January and February at this site. This field was surrounded by 10 foot (3-meter) fencing to preclude deer foraging. Preparation and planting followed Stearns and Fenster (2016a), which includes removing preexisting vegetation and raking the soil. Seedlings were planted 10 cm apart at the 2-to 4-leaf stage. There were 60 plants for each subline of both mutant lines and premutation founders (with few exceptions, Table 1) for a total of 5,520 plants. For all founders, we planted 120 seedlings for the control. For Columbia, we planted an additional 2,400 mutated seedlings (20 mutant lines). For Cape Verde (CV), we planted 1920 mutated seedlings (16 mutant lines).

**Table 1.**
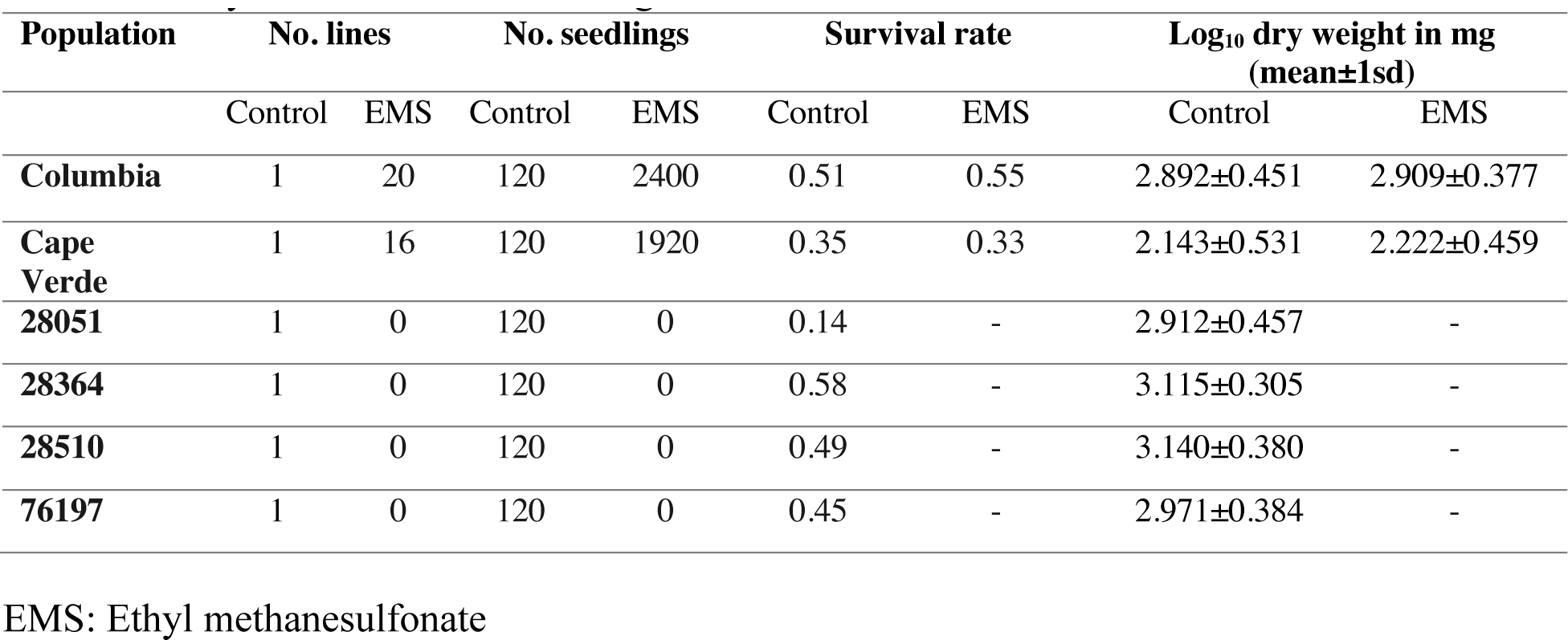
Summary statistics of mutation experiments for the two founders (Columbia and Cape Verde) subjected to both the mutation and control treatment and the four founders (28051, 28364, 28510, 76197) subjected to only the control treatment. Log10 mg dry weight was calculated only for the survived seedlings.

| Population | No. lines |  | No. seedlings |  | Survival rate |  | Log<sub>10</sub> dry weight in mg<br>(mean±1sd) |  |
| --- | --- | --- | --- | --- | --- | --- | --- | --- |
|  | Control | EMS | Control | EMS | Control | EMS | Control | EMS |
| **Columbia** | 1 | 20 | 120 | 2400 | 0.51 | 0.55 | 2.892±0.451 | 2.909±0.377 |
| **Cape Verde** | 1 | 16 | 120 | 1920 | 0.35 | 0.33 | 2.143±0.531 | 2.222±0.459 |
| **28051** | 1 | 0 | 120 | 0 | 0.14 | - | 2.912±0.457 | - |
| **28364** | 1 | 0 | 120 | 0 | 0.58 | - | 3.115±0.305 | - |
| **28510** | 1 | 0 | 120 | 0 | 0.49 | - | 3.140±0.380 | - |
| **76197** | 1 | 0 | 120 | 0 | 0.45 | - | 2.971±0.384 | - |

EMS: Ethyl methanesulfonate

The plots were initially watered for a week to help with establishment but otherwise exposed to natural conditions. Plants were harvested in May of 2015 when all plants had senesced. Plants were then dried in heat chambers. Above ground biomass has been shown to be strongly correlated to fruit number in earlier experiments with these lines (Shaw et al. 2000; Stearns and Fenster 2016b,a; Weng et al. 2021). Therefore, this biomass measure was used as an estimate of fitness. Our method of quantifying fitness, total seed set for an annual plant, is a standard approach in plant ecological genetics (Schemske 1984; Briggs and Walters 2016). Measuring seed production is a robust estimate of reproductive success since *A. thaliana* is highly selfing (e.g., Stenøien et al. 2005). Thus, we were able to efficiently quantify the distribution of fitness effects of new mutations measured in an environment approximating natural selection. Many previous experiments to study the fitness effects of new mutations using microorganisms are constrained to laboratory environments (e.g. Baer et al. 2006), but do have the advantage of measuring fitness as population growth rates, which is an arguably more direct estimate of fitness.

### Bayesian inference of genetic and nongenetic parameters

Our main interest in this paper is to estimate the parameters for the distribution of fitness effects (DFE) of mutations from the phenotypic measurements. To do this, we first decompose the measurement in M4 (Figure 1) as a sum of genetic and nongenetic terms:

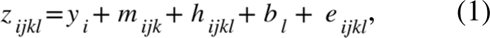

Where *z_ijkl_* is the measurement of founder *i*, line *j*, subline *k*, and individual *l*. *y_i_* is the genetic value of the founder *i*. *m_ijk_* is the subline specific maternal effect. *h_ijkl_* is the deviation in the offspring’s genetic value from the founder that contains the effects of mutations that fixed in the common ancestor of line *j*, as well as mutations that subsequently fixed in subline *k*. *b_l_* is the block effect associated with individual *l*. *e_ijkl_* is the residual due to phenotypic and measurement noise.

Here we assume that the number of mutations contributing to *h_ijkl_* follows a Poisson distribution with the mean given by the mutation rate calculated in the previous section. We further assume that there is no epistasis and dominance. Additivity of mutations on fitness in *A. thaliana* has been documented for later life-history traits (Shaw and Chang 2006). We also expect that 87.5% of the mutations will be homozygous because of the 3 generations of selfing to create the mutant lines, so that dominance will likely account for a small proportion of the observed genetic variation even when present. Therefore, *h_ijkl_* is equal to the sum of a Poisson-distributed number of independent mutational effects drawn from the same DFE. Our experimental design with two sublines allows us to estimate the contribution of maternal effects *m_ijk_* to total variance, which could otherwise inflate our estimate of mutational variance if unaccounted for. However, this design also introduces some challenges to the statistical analyses, in that the mean values of total mutational effects of the two sublines are now correlated, due to their shared mutations fixed in the M2 generation as well as mutations that were still segregating in M2 (see Figure 1, Supplemental Figure 1). In the Supplement, we detail the decomposition of *h_ijkl_* into effects of mutations fixed at different generations, which allow us to explicitly model the correlation between sublines.

In order to estimate the DFE, as well as the distributions of maternal effects and residual noise, we use a hierarchical Bayesian approach to model the random effects (*m_jk_*, *h_ijkl_*, *b_l_*, *e_ijkl_*) in Eq. 1. Specifically, we specify that a random genetic or nongenetic effect is drawn from a probability distribution (for instance the DFE for effects of individual mutations), whose parameters (e.g. the mean and standard deviation of the DFE) in turn follow a prior distribution specified by their respective hyperparameters. This allows us to not only calculate the point estimate for the parameters of interests, but also their entire posterior distribution.

To construct a prior distribution for the DFE, previous papers have required the DFE be within a designated family of probability distributions (e.g. Gamma distribution, Keightley 1994) and impose prior distributions over the distribution parameters. This approach relies on strong a priori assumptions on the shape of the DFE, which could potentially introduce biases into the results. Here we realize that we can circumvent this problem by taking advantage of the relatively large number of mutations introduced by EMS. Specifically, note that *h_ijkl_* is a sum of a Poisson distributed number of i.i.d mutational effects. We can accordingly use the Central Limit Theorem to show that when the number of mutations per offspring is large, the distribution of *h_ijkl_* can be well approximated by a normal distribution (see the Supplement for proof), regardless of the shape of the DFE. This allows us to take a nonparametric approach to estimate the mean *μ* and standard deviation *σ* of the DFE without restricting to certain distribution families.

Additionally, we specify the random maternal effect *m_jk_* block effect *b_l_*, and the noise term *e_ijkl_* to follow zero mean normal distributions, whose standard deviations are drawn from uninformative priors. The founder’s genetic value *y_i_* is treated as a fixed effect and sampled from an uninformative zero-mean normal distribution with fixed standard deviation.

One problem in directly using Eq. 1 is that our fitness measurements contain many zeros since many seedlings or juvenile plants (prior to reproduction) died before measurements could be made. To solve this problem, we re-formulate our observation *z_ijkl_* to follow a mixture distribution. We do this by modeling the probability of surviving to the time of measurement for every plant *ijkl* as a logistic function of the sum of its genetic effects and maternal effects. That is

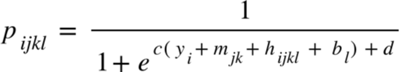

Here *c* and *d* are parameters that determine the shape of the logistic function and are specified to follow uninformative prior distributions. With this modification, *z_ijkl_* is equal to zero with Probability 1 − *p_ijkl_*. In the event that the seedling survived to measurement with probability *p_ijkl_*, *z_ijkl_* is nonzero and follows a normal distribution with mean equal to *y_i_* + *m_jk_* + *h_ijkl_* + *b_l_*, and variance equal to the variance of the noise term *e_ijkl_*.

Ideally, we would like to perform Monte Carlo sampling for both the model parameters and individual random effects. However, this is not practical due to the complexity of our model. Therefore, to estimate the posterior distribution for the model parameters, we use the variational Bayesian method (Fox and Roberts 2012), which approximates the complex intractable posterior of the parameter *θ* given the data X, *P*(*θ*∣*X*) with a simple factorized distribution *Q*(*θ*), by minimizing the log odds between *Q*(*θ*) and the joint distribution *P*(*θ*, *X*) expected under *Q*(*θ*), which provides a lower bound for Kullback–Leibler divergence between *Q*(*θ*) and *P*(*θ*∣*X*). We performed variational inference for our model using the Python package ’pymc3’ (Salvatier et al. 2016).

## Results

### Summary statistics

For the control lines across all six accessions, the overall survival rate is 0.42, while the mean±1std dry weight in log_10_ mg for the survived seedlings is 2.86 ±0.45.

The overall survival rate calculated across the two accessions that received the EMS mutagenesis treatment (Columbia and Cape Verde) is 0.45, while the mean±1std dry weight in log_10_ mg for the survived seedlings is 2.60±0.52.

For Columbia (COL), the overall survival rate is 0.544. Specifically, the survival rate for the mutated seedlings was 0.546, compared with the survival rate of 0.508 for the control. The log_10_ mg dry weight of all survived seedlings is 2.909 ±0.377 for the mutation lines, and 2.891±0.451 for the control lines (See Figure 2, Table 1).

**Figure 2.**
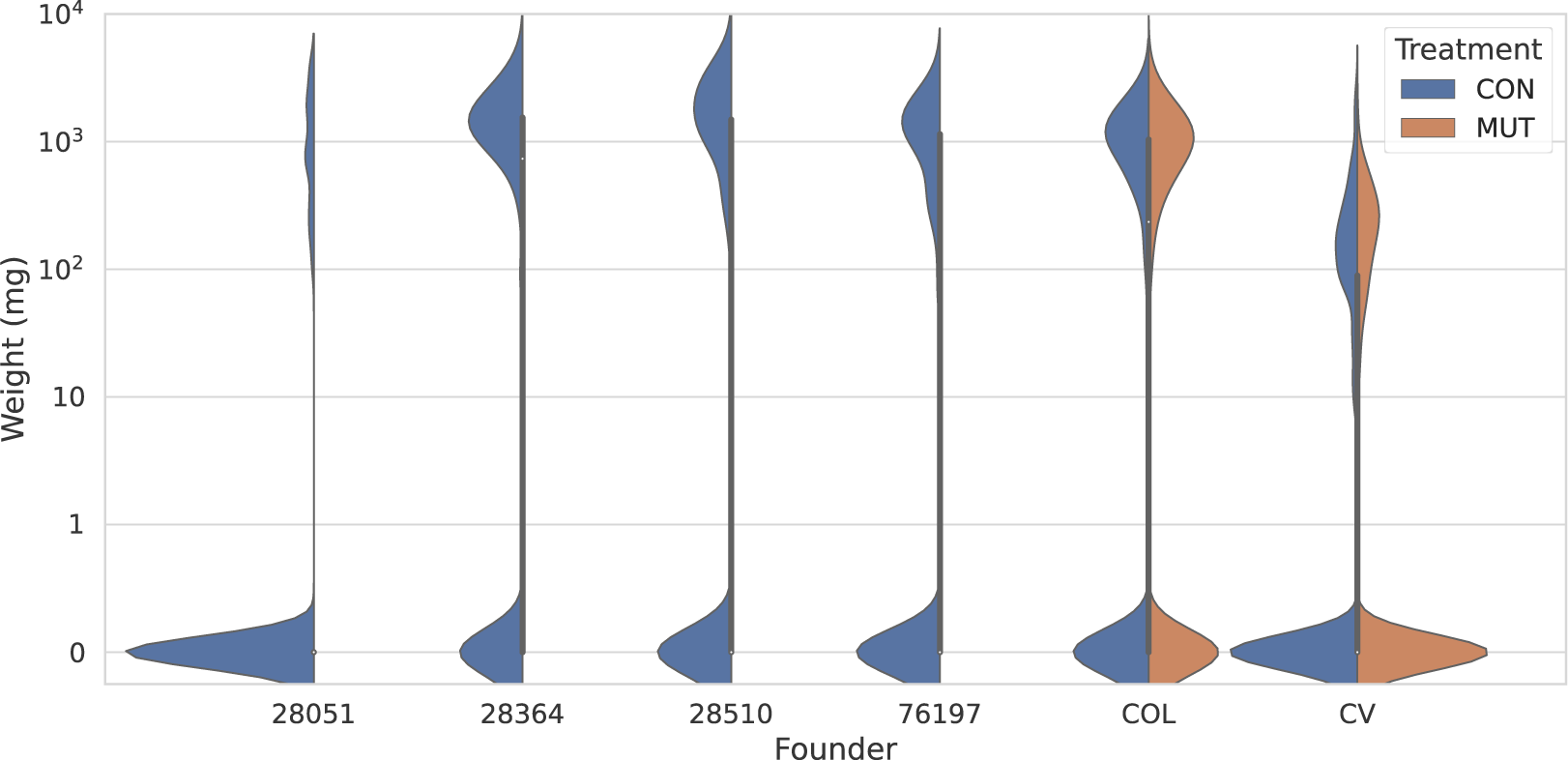
Distribution of dry weight of seedlings for the six founders. For the two founders (Columbia and Cape Verde) that received mutation accumulation treatment, the distributions are separately plotted for the control and the mutation lines. Note that the distributions are bimodal. The probability concentered at zero corresponds to seedlings that failed to germinate (i.e. dry weight=0). Density estimation was performed using the kernel density estimation method with kernel bandwidth=0.09, and grid size=100. CON: control. MUT: EMS mutation treatment. COL: Columbia. CV: Cape Verde.

For Cape Verde (CV), we observed lower survival rate as well as dry weight than the COL line. Specifically, overall survival rate across both mutation and control treatment is 0.333 (seedling survival rate is 0.332 across all COL mutation lines, and 0.350 for the control line). The log_10_ dry weight of the survived seedlings is 2.222±0.459 for the mutation lines, and 2.143+0.531 for the control line (See Figure 2, Table 1).

### Posterior estimates of distribution of fitness effects of mutations

Note that since we used a hierarchical Bayesian approach, the parameters of interests such as the mean and the standard deviation of DFE have their respective posterior means, variances, and credible intervals. All quantities reported are in units log_10_ mg.

Overall, our point estimations of the block effects are small, with mean±1sd = −0.02±0.02 across all 12 blocks. In Fig 3 and Table 2, we present the posterior distribution of model parameters of the DFE, as well as the residual noise distribution and founder genetic values. The difference in seedling survival rates and the distributions of dry weight between CV and COL can be largely explained by the estimate that the genetic value (y) of CV founder before the mutation treatment is much lower than COL and other lines. Specifically, the posterior mean ± 1sd of genetic value for CV founder is y=2.164±0.017, compared with y=2.886±0.009 for the COL founder (Figure 3A, Table 2).

**Figure 3.**
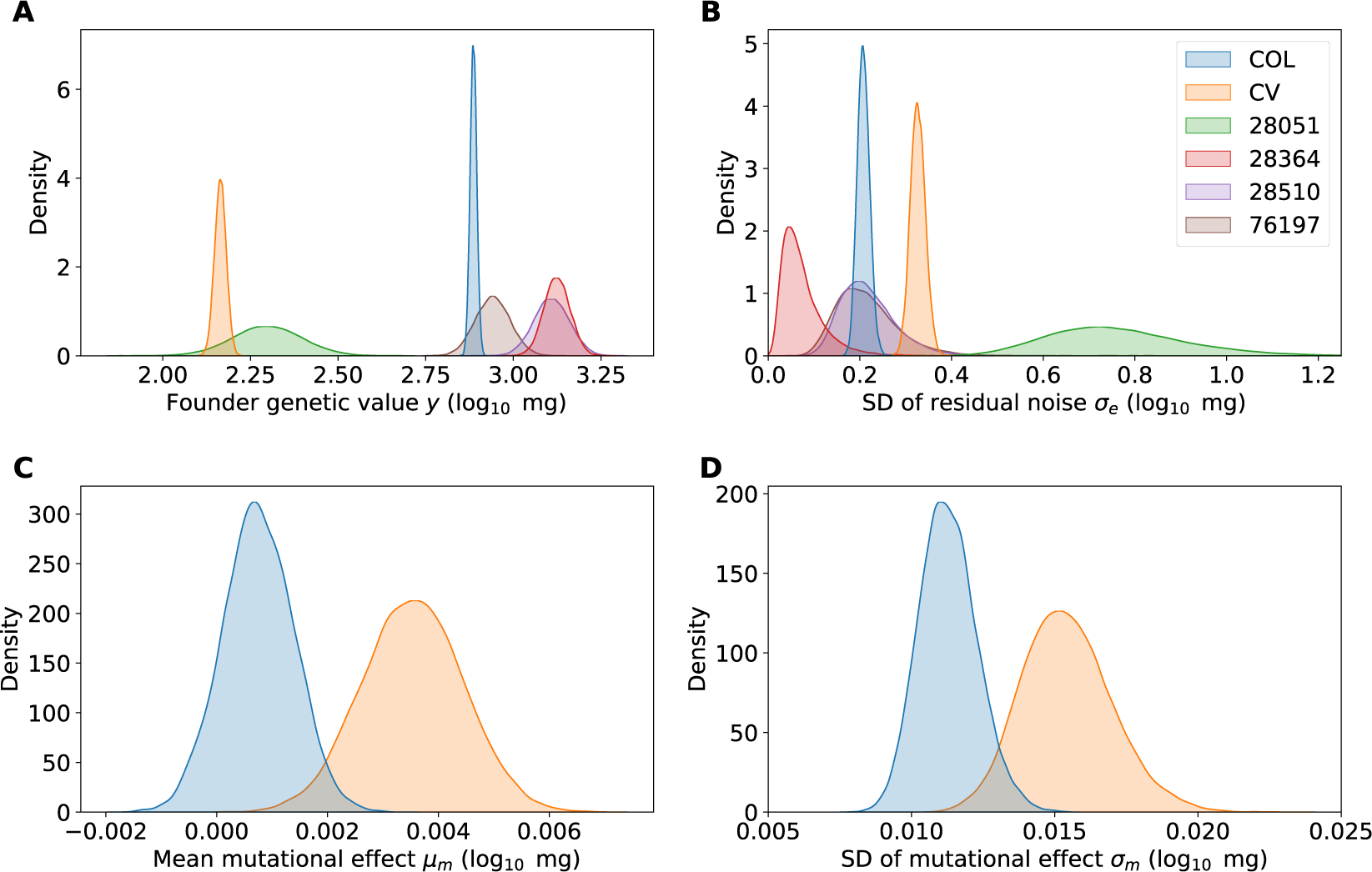
Posterior distributions for various parameters inferred by fitting a Bayesian hierarchical model to the data. (A) founder’s genetic value y; (B) standard deviation of the residual noise σ_e_; (C) mean of the distribution of the fitness effect (DFE) of mutations μ_m_; (D) standard deviation of the DFE, σ_m_. COL: Columbia. CV: Cape Verde.

**Table 2.** Parameters of the distribution of fitness effects (DFE). The numbers correspond to the posterior means of our Bayesian model. Note that the DFE parameters were only calculated for the two founders (Columbia and Cape Verde) that received the EMS mutation treatment.

| Population | y | $\mu_m$ | $\sigma_m$ | $\sigma_e$ | $h_m^2$ | $CV_m$ |
| --- | --- | --- | --- | --- | --- | --- |
| **Columbia** | 2.886 | 0.001 | 0.011 | 0.208 | $0.447 \times 10^{-3}$ | 0.566 |
| **Cape Verde** | 2.164 | 0.004 | 0.015 | 0.327 | $0.337 \times 10^{-3}$ | 0.775 |
| **28051** | 2.298 |  |  | 0.762 |  |  |
| **28364** | 3.125 |  |  | 0.074 |  |  |
| **28510** | 3.109 |  |  | 0.218 |  |  |
| **76197** | 2.941 |  |  | 0.215 |  |  |

y: genetic value of the founder. $\mu_m$ : Mean of the DFE. $\sigma_m$ : standard deviation of the DFE. $\sigma_e$ : standard deviation of the residual noise. $h_m^2$ : mutational heritability. $h_m^2 = 0.16 \sigma_m^2 / \sigma_e^2$ . $\mu_m$ , $\sigma_m$ , and $\sigma_e$ are in units $\log_{10}$ mg.

The results for the three founders without mutation treatment are y=2.298±0.099 for accession 28051, y=3.125±0.037 for 28364, y=3.109±0.05 for 28510, and y=2.941±0.05 for 76197 (Figure 3A, Table 2). Note that the posterior distribution for genetic value is much narrower for CV and COL due to their larger sample sizes (control + mutation) compared with the other three founders (n=120 only for the control group).

The posterior distribution of the standard deviation of residual noise (σ_E_) is presented in Figure 3B. CV exhibits higher residual noise variance than COL: σ_E_=0.327±0.016 for CV; σ_E_=0.208±0.013 for COL (Table 2). Here again the posterior distributions of the three lines without mutation treatment are much wider, due to the low sample size.

The posterior distribution of the mean of the DFE (μ_m_) for CV and COL is presented in Figure 3C. The posterior distribution for the mean mutational effect of COL is close to zero, with μ_m_ = 0.00075±0.00064, and a 95% credible interval that intersects 0. In contrast, for CV, μ_m_ =0.00354±0.00091, with 99% credible interval not intersecting zero (Table 2). Therefore, mutations in the lower fitness CV background overall have slightly positive effects. This contrasts with the relatively higher performing COL founder, where a mutation has equal probability of being beneficial and deleterious.

Finally, in Figure 3D, we examine the posterior distribution of the standard deviation of the DFE (σ_m_) of CV and COL. For the Columbia accession, which appears to be more adapted to the experimental field site, σ_m_ =0.011±0.001, whereas CV exhibits higher variation in its mutational effects, with σ_m_ =0.015±0.002 (Table 2).

We also calculated the per-generation increase in heritability due to mutation (mutational heritability, h^2^_m_) as h^2^_m_ = V_m_/V_e_ (Houle et al. 1996), where we first set V_e_ = σ_e_^2^. The new variance due to mutation (V_m_) is equal to the variance of the effect of single mutations (σ_m_^2^) times the expected number of mutations per generation in the absence of EMS (diploid genomic rate of mutation rate affecting biomass and fitness = 0.16 per generation, Weng et al. 2019). Using this equation, we found the mutational heritability for Col and CV accessions extrapolated from EMS to new mutations is h^2^_m_ = 0.447 ×10^-3^, and h^2^_m_ = 0.337 × 10^-3^ (Table 2), respectively. Assuming that the log_10_ function is well approximated by a linear function close to the founder mean, these values are largely consistent with the previously observed h^2^_m_ for biomass in *A. thaliana* mutation lines (Weng et al. 2021). In addition, we can also calculate the variance in genetic values among the six accessions (namely V_g_) using values in Table 1 and found V_g_ = 0.174, which is 8530 and 4563 times higher than the V_m_ of COL and CV lines, respectively.

## Discussion

### Distribution of mutational effects

Using chemically induced mutations we confirm the results of prior *A. thaliana* MA line studies based on natural new mutations that demonstrated that the frequency of beneficial mutations is much greater than predicted by the prior literature (Lynch and Walsh 1998). Our study, with mutational lines from two founders is consistent with 25 previous founder in field (Rutter et al. 2010, 2018; Roles et al. 2016; Weng et al. 2021) and greenhouse (Shaw et al. 2002; Rutter et al. 2010; Stearns and Fenster 2016b) *A. thaliana* studies that quantified a higher frequency of beneficial mutations than previously expected. Our results are also consistent with several studies that have quantified mutation effects on *A. thaliana* performance in laboratory conditions, where the mutagen is a single T-DNA insert (Rutter et al. 2017; Murren et al. 2019). Spoelhof et al. (2021) found EMS induced mutations in *A. thaliana* to have uniformly deleterious effects on performance. However, they used a higher dosage and did not transfer the mutations through several generations (as we did). Transferring mutations across generations eliminates lethal and highly deleterious mutations. A number of other studies with other organisms have also documented a higher than expected frequency of beneficial mutations since the initial observation that a high proportion of mutations found in *A. thaliana* MA lines may be beneficial (Perfeito et al. 2007, 2014; Silander et al. 2007; Hall et al. 2008; Latta IV et al. 2013; Böndel et al. 2019, 2022), challenging the dogma that mutations are nearly uniformly deleterious (Keightley and Lynch 2003).

There are at least two explanations for why we recover beneficial mutations with such high frequency in *A. thaliana*. First, if the DFE is highly leptokurtic, as observed by Böndel et al. (2022), then mutations will mostly have very small effect on fitness, but over time a line or lineage will eventually incorporate a highly deleterious mutation that will cause its extinction. Under this scenario, our dosage of EMS, equivalent to 150 generations of mutation accumulation was not high enough to simulate a long enough period to capture those very deleterious mutations. The second explanation is the genotype × environment (G×E) argument that genotypes are rarely at their adaptive optimum, and if mutations have relatively small effect on fitness, then there will be a relatively high frequency of beneficial mutations (Martin and Lenormand 2015).

Our results are consistent with some theoretical and empirical predictions. Since the four non-Cape Verde and Columbia accessions performed better than either Cape Verde or Columbia, we can assume that neither Cape Verde nor Columbia accessions were well adapted to our specific field environment. With a uniform fitness surface, we expect a higher proportion of beneficial mutations if the genotype is far from its adaptive optimum (Fisher 1930), as we have documented here. The Cape Verde accession has a higher frequency of beneficial mutations than Columbia, which is also consistent with the adaptive optimum argument since Cape Verde has low fitness compared to other accessions in the study area (Rutter and Fenster 2007; Stearns and Fenster 2016a). Furthermore, *A. thaliana* populations at the edge of the species distribution harbor more maladaptive loci, likely due to drift and population bottlenecks (Oakley et al. 2023). Since the Cape Verde accession is an isolated population, far from the main range of *A. thaliana* in Europe, it is possible that it has a relatively higher genetic load, setting the stage for beneficial compensatory mutations as observed in other mutation studies (e.g., Burch and Chao 2000).

However, if the DFE of new mutations is leptokurtic (Böndel et al. 2019, 2022), we might have expected to recover severe performance reducing mutations, but we did not. It is possible that our method of mutation induction may have led to a form of somatic selection against deleterious mutations; however, previous results suggest that somatic selection acting on new mutations is not significant (Monroe et al. 2022). It is important to note that we are unable to reconstruct the shape of the DFE of individual mutations, only the average effect of mutations. This may obscure important details of a stepwise adaptive “walk.” Further, the shape of the mutational DFE has important consequences on evolutionary process from the validity of the molecular clock to how quickly populations can respond to selection via mutations (Böndel et al. 2022). Therefore, future work needs to address both the proportion of new mutations that can contribute to selection response, as well as the exact shape of the DFE.

### Mutational input to genetic variation

While it is clear that standing genetic variation contributes to selection response, it is difficult to quantify the relative contributions of new mutations to this dynamic (Barrett and Schluter 2008). Our results, based on the production of new mutations that were tested in semi-natural environments demonstrate that new mutations are potentially beneficial often enough and of large enough effect size, such that when combined with high mutation rate, they may contribute significantly to rapid local adaptation under natural conditions.

The mean mutational heritability for Columbia and Cape Verde is 0.392 × 10^-3^, suggesting that mutation alone could generate typical h^2^ of 0.5 (Mousseau and Roff 1987; Roff 2012) in about 1300 generations. Thus, populations can respond to selection pressures via new mutations in a relatively limited number of generations, consistent with previous laboratory investigations of mutational input to *D. melanogaster* selection response (Mackay et al. 1992, 1994).

The variance in genetic value among the six accessions (V_g_=0.174) is 8,530 and 4,563 times higher than the genetic variance generated by mutations alone (V_m_, see Table 2) of COL and CV lines, suggesting that it would take roughly 4,500-8,500 generations of mutation accumulation alone to produce the observed phenotypic variance among the six accessions at our study site due to genetic contributions (Mousseau and Roff 1987; Roff 2012). Furthermore, our results also allow us to estimate the time scale on which Cape Verde is able invade the fitness space of Columbia. Specifically, assuming zero mean and constant standard deviation over future generations, the number of generations for the distribution of fitness of Cape Verde to contain the mean genetic value of Columbia within 2 standard deviations from its own genetic value, e.g. (y_COL_ - y_CV)_^2^/(4 × V_M, Cape Verde_) = 3417 generations, using values given in Table 2. Thus, low fitness Cape Verde ecotype is able to occupy the “adaptive space” of a higher fitness genotype in ∼3400 generations to become equivalently adapted to the habitat, which is on the same order of magnitude as the number of generations needed for establishing the genetic variance among all ecotypes as mentioned above. Overall, we conclude that our results contribute to an increasing number of studies (e.g., Izutsu and Lenski 2022) that demonstrate that new mutations can play a significant role in the evolutionary process at the population level.

We were also able to identify the contribution of “phenotypic noise” (phenotypic variation that is not due to genetic variation). We find that there is greater phenotypic noise in the lower fitness ecotypes than the higher ecotypes. This fits predictions from recent theoretical work (Rocabert et al. 2020) and corroborates recent experimental work that found that noisy phenotypes can be more beneficial under stressful conditions (Samhita et al. 2021). Although this variation is not heritable and does not contribute directly to adaptation, it may play a role in preventing population extinction during the “waiting time” for additional adaptive mutations (the “look-ahead” effect of Whitehead et al. 2008). Therefore, although phenotypic noise was not a focus of this study, this intriguing result deserves further investigation.

### Novelty and limits of the Bayesian analysis

Our experimental design and Bayesian analysis utilize several novel approaches to estimating the DFE in field environments. Most notably, our inference method is nonparametric, in that it does not assume the shape of the DFE and instead directly characterizes the DFE by estimating the mean and variance of mutational effects. This is thanks to the large number of mutations introduced by the EMS treatment, which allowed us to use the Central Limit Theorem to approximate the joint genetic effect of a relatively large, Poisson-distributed number of mutations. Classical approaches of modeling the DFE usually assume that the DFE follows certain distributions, e.g., the Gamma distribution. However, it has been shown that departures of the true DFE from the assumed distribution (especially if the DFE is bimodal or multimodal) can lead to erroneous estimations of the DFE parameters (Kousathanas and Keightley 2013). Therefore, hypothesis-free, nonparametric models may prove to be more desirable approaches since they are likely less prone to biases caused by model misspecifications.

Furthermore, we took advantage of the flexibility of Bayesian models to account for several complexities in the data that cannot be fully captured by Frequentist approaches. These include explicitly modeling the covariance between sublines due to shared segregating mutations, as well as using a logistic function to model the survival of seedlings as a function of the underlying genetic value, which allowed us to jointly model the zero-inflated fitness data in a principled way. The general Bayesian modeling approach used in this paper may be relevant for other studies of DFEs and may allow researchers to design more complex mutagenesis and MA experiments to characterize finer-scale properties of the genetic effects of mutations.

One potential drawback of our inference procedure is that due to the complex data structure, we used the variational inference procedure, instead of unbiased sampling methods such as MCMC. The variational inference method approximates the posterior distribution using simple, factorized distributions to allow fast inference (Fox and Roberts 2012). However, this may potentially introduce bias into our estimates. Future work on efficient computational methods may be needed to fit data generated from complex mutagenesis experiments.

### Summary

Using chemically induced mutagenesis and Bayesian nonparametric modeling, we estimated the distribution of fitness effects of mutations in field environments for two ecotypes that are differentially adapted to the local environment. We found that large proportions of mutations are beneficial for both ecotypes and that the ill-adapted ecotype showed higher variance in the effect of mutations as well as a slight bias towards beneficial mutations in its DFE. Our results suggest that beneficial mutations when combined with high mutation rate and potentially high phenotypic noise can contribute to local adaptation on ecological time scales.

## Author Contributions

FWS and CBF designed the experiment. FWS and CBF conducted the field experiment. JZ performed the Bayesian analyses. FWS, JZ, and CBF wrote the paper.

## Acknowledgements

This work was supported by NSF DEB 1257902 to C. Fenster, and College of Liberal Arts and Sciences at the University of Florida. We thank Charles Baer for valuable feedbacks for the interpreting the data.

## Data and Code availability

Fitness data and Python code for performing the analyses and recreating all figures can be found at Github repository: github.com/juannanzhou/Arabidopsis_DFE.

