## Supplemental methods for "Scaling the fitness effects of mutations with respect to differentially adapted *Arabidopsis thaliana* accessions under natural conditions"

### Model description

We fit the following model to our fitness data:

$$y_{ijkl} = y_i + m_{ijk} + h_{ijkl} + b_l \tag{1}$$

$y_{ijkl}$ is the sum of genetic, maternal, and block effects for of founder $i$, line $j$, subline $k$, and individual $l$. Specifically, $y_i$ is the genetic value of the founder $i$. $m_{ijk}$ is the subline specific maternal effect. $b_l$ is the block effect. $h_{ijkl}$ is the individual specific genetic deviation from the genetic value of the founder due to de novo mutations.

We assume that subline-specific maternal effects are drawn from a normal distribution:

$$m_{ijk} \sim \text{Normal}(0, \gamma^2), \tag{2}$$

where $\gamma$ is a hyperparameter that follows a flat prior

$$\gamma \sim \text{HalfNormal}(10) \tag{3}$$

For each founder, we assume that genetic value follows an uninformative normal distribution $y_i \sim \text{Normal}(6.2, 10)$, where 6.2 is equal to the grand mean of fitness of all measured seedlings. We next define founder specific distribution of fitness effects (DFE) of mutations, such that for founder $i$, the mean $\mu_{m,i}$ and standard deviation $\sigma_{m,i}$ of the DFE both follow flat priors:

$$\mu_{m,i} \sim \text{Normal}(0, 10) \text{ and } \sigma_{m,i} \sim \text{HalfNormal}(10). \tag{4}$$

Due to our experimental design where two sublines were established in generation 3 (M3, Main Text Figure 1), the fitness values of the seedlings of the two sublines are non-independent due to shared mutations in their common ancestor in M2. Here we provide detailed derivation for the distribution of $h_{ijkl}$. Specifically, we decompose $h_{ijkl}$ as

$$h_{ijkl} = f_{ij} + s_{ijk} + r_{ijkl}. \tag{5}$$

Here $f_{ij}$ is the genetic effect of mutations fixed in M2 for founder $i$ and line $j$. This effect is shared by both sublines. $s_{ijk}$ is deviation of genetic values of the ancestor in M3 due to segregating sites in M2. $r_{ijkl}$ is the deviation in genetic values for individual $ijkl$ due to segregating mutations in M3.

We begin our derivation at generation 0. Here the genome is homozygotes for all sites. In the next generation, M1, there are a number of heterozygous sites due to de novo mutations. The number follows a Poisson distribution with mean $\lambda = 25$, which is empirically derived from previous experimental data (see Main Text for details).

In M2, on average a quarter of the segregating sites are fixed for the mutant alleles. Let this number be $N$. Then $N$ follows a Poisson distribution with mean $\lambda/4$. These mutations contribute to the value $f_{ij}$, such that

$$f_{ij} = \sum_{m=1}^{N} 2e_m, \tag{6}$$

where $e_m$ is the effect of a single mutation that is sampled from the line-specific DFE. The factor 2 corresponds to the fact that the loci under consideration are homozygous for the mutant alleles. This is a compound Poisson distribution. Based on the Central Limit Theorem, this distribution asymptotically approximates the normal distribution when the mean of $N$ is large. Thus here we use a normal distribution to approximate the distribution of $f_{ij}$. This allows us to avoid directly sampling the effects $e_m$, which can substantially reduce the computational burden. Using the law of total variance [1],

$$\text{Var}(f_{ij}) = \mathbb{E}_M[\text{Var}(f_{ij}|N)] + \text{Var}(\mathbb{E}[f_{ij}|N]), \tag{7}$$

we can derive the mean and variance of $f_{ij}$, such that its distribution can be approximated with a normal distribution with the same first two moments

$$f_{ij} \sim \text{Normal}(\frac{\lambda}{2}\mu_{m,i}, \lambda \times (\sigma^2_{m,i} + \mu^2_{m,i})) \tag{8}$$

In addition to the fixed mutations in M2 that give rise to effect $f_{ij}$, there are also on average $P = \lambda/2$ mutations that were still segregating in this generation. This mutations will be sorted into the founders of the two sublines in M3, giving rise to the subline specific deviation $s_{ijk}$. This value again follows a compound Poisson distribution

$$s_{ijk} = \sum_{m=1}^{P} n_m \times e_m. \tag{9}$$

Here $P$ follows a Poisson distribution with mean $\lambda/2$. $n_m = 0, 1, 2$ corresponds to the dosage of the mutant allele on the segregating site $m$, which follows a categrocial distribution

$$n_m \sim \text{Categorical}(\begin{bmatrix}0\\1\\2\end{bmatrix}, \begin{bmatrix}1/4\\1/2\\1/4\end{bmatrix}), \tag{10}$$

which has a mean 1. For the effects of mutations at a single locus $n_m \times e_m$, it is easy to see

$$\mathbb{E}[n_m \times e_m] = \mu_{m,i}. \tag{11}$$

It has a variance

$$\begin{aligned}
\text{Var}(n_m \times e_m) &= \mathbb{E}[(n_m \times e_m)^2] - \mathbb{E}[(n_m \times e_m)]^2 && (12)\\
&= \mathbb{E}[n_m^2]\mathbb{E}[e_m^2] - \mu^2_{m,i} && (13)\\
&= 1.5 \times (\sigma^2_{m,i} + \mu^2_{m,i}) - \mu^2_{m,i} && (14)\\
&= 1.5\sigma^2_{m,i} + 0.5\mu^2_{m,i} && (15)
\end{aligned}$$

Due to the fact that the $P$ segregating sites are common among the two sublines, $s_{ijk}$ for $k = 1$ and 2 are not independent. Here we use a bivariate normal distribution to approximate their joint distribution. Using the law of total covariance [3]

$$\text{Cov}(s_{ij1}, s_{ij2}) = \mathbb{E}_P[\text{Cov}(s_{ij1}, s_{ij2}|P)] + \text{Cov}(\mathbb{E}[s_{ij1}|P], \mathbb{E}[s_{ij2}|P]), \tag{16}$$

and Eqs. 11, 12, we can show that

$$\begin{bmatrix}s_{ij1}\\s_{ij2}\end{bmatrix} \sim \text{Normal}\left(\frac{1}{2}\begin{bmatrix}\lambda\mu_{m,i}\\\lambda\mu_{m,i}\end{bmatrix}, \frac{1}{2}\begin{bmatrix}1.5(\sigma_i^2 + \mu^2_{m,i})\lambda & (\sigma_i^2 + \mu^2_{m,i})\lambda\\(\sigma_i^2 + \mu^2_{m,i})\lambda & 1.5(\sigma_i^2 + \mu^2_{m,i})\lambda\end{bmatrix}\right) \tag{17}$$

provides an approximation for the joint distribution of the subline specific genetic deviations.

Finally, we have the value $r_{ijkl}$, which corresponds to deviation in genetic values for individual $ijkl$ due to segregating mutations in M3. Due to the relatively low number of remaining segregating sites (mean $= \frac{1}{4}\lambda$), here we approximate the distribution of $r_{ijkl}$ as independent, identical Gaussian distributions such that

$$r_{ijkl} \sim \text{Normal}(\frac{\lambda}{4}\mu_{m,i}, \frac{1}{4}\lambda \times 1.5(\sigma_i^2 + \mu^2_{m,i})), \tag{18}$$

which is derived again using the law of total variance and Eqs. 11, 12.

To accomodate the many zeros in our dataset caused by failure of gemination and death of seedlings, we assume that that the probability of an individual survives to the time of measurement $p_{ijkl}$ follows a logistic function depending on the joint genetic, maternal, and block effect

$$p_{ijkl} = \text{Logistic}(c * z_{kijl} + d), \tag{19}$$

where $c$ and $d$ specify the shape of the logistic function and follow uninformative priors. Thus, the fitness measurement follows a mixture distribution

$$z_{ijkl} = (1 - \text{Bernoulli}(p_{ijkl})) \times 0 + \text{Bernoulli}(p_{ijkl}) \times (y_{ijkl} + \varepsilon_{ijkl}) \tag{20}$$

$\varepsilon_{ijkl}$ is the measurement noise. We assume that the residual noise follows a founder specific normal distribution:

$$\varepsilon_{ijkl} \sim \text{Normal}(0, \sigma^2_{e,i}), \tag{21}$$

$$\sigma_{e,i} \sim \text{HalfNormal}(10). \tag{22}$$

For convenience, we summarize the distribution of all random variables under our hierarchical model below.

| Variable | Distribution |
|---|---|
| SD of maternal effect | $\gamma \sim \text{HalfNormal}(10)$ |
| Block effect | $b_l \sim \text{Normal}(0, 10)$ |
| **Founder**: $i$ | |
| SD of residual noise | $\sigma_{e,i} \sim \text{HalfNormal}(10)$ |
| Founder's genetic value | $y_i \sim \text{Normal}(6.2, 10)$ |
| Mean of mutational effect of founder | $\mu_{m,i} \sim \text{Normal}(0, 10)$ |
| SD of mutational effect of founder | $\sigma_{m,i} \sim \text{HalfNormal}(10)$ |
| **Line**: $j$ | |
| Fixed mutational effects in M2 | $f_{ij} \sim \text{Normal}(\frac{\lambda}{2}\mu_{m,i}, \lambda \times (\sigma_{m,i}^2 + \mu_{m,i}^2))$ |
| **Subline** $k$ | |
| Maternal effect | $m_{ijk} \sim \text{Normal}(0, \gamma^2)$ |
| Subline specific genetic effects | $\begin{bmatrix} s_{ij1} \\ s_{ij2} \end{bmatrix} \sim \text{Normal}\left(\frac{1}{2}\begin{bmatrix} \lambda\mu_{m,i} \\ \lambda\mu_{m,i} \end{bmatrix}, \frac{1}{2}\begin{bmatrix} 1.5(\sigma_i^2 + \mu_{m,i}^2)\lambda & (\sigma_i^2 + \mu_{m,i}^2)\lambda \\ (\sigma_i^2 + \mu_{m,i}^2)\lambda & 1.5(\sigma_i^2 + \mu_{m,i}^2)\lambda \end{bmatrix}\right)$ |
| **Individual**: $l$ | |
| Individual specific genetic effects | $r_{ijkl} \sim \text{Normal}(\frac{\lambda}{4}\mu_{m,i}, \frac{1}{4}\lambda \times 1.5(\sigma_i^2 + \mu_{m,i}^2))$ |
| Sum of genetic, maternal, and block effects | $y_{ijkl} = y_i + m_{ijk} + f_{ij} + s_{ijk} + r_{ijkl} + b_l$ |
| Probability of survival of subline | $p_{ijkl} = \text{Logistic}(c * y_{ijkl} + d)$ |
| Residual noise | $\varepsilon_{ijkl} \sim \text{Normal}(0, \sigma_e^2)$ |
| The observed fitness | $z_{ijkl} = (1 - \text{Bernoulli}(p_{ijkl})) \times 0 + \text{Bernoulli}(p_{ijkl}) \times (y_{ijkl} + \varepsilon_{ijkl})$ |

The likelihood function is then given by

$$\prod_{i,j,k,l} p(z_{ijkl}|y_{ijkl}, \sigma_{\varepsilon,i}) \times p(r_{ijkl}|\mu_{m,i}, \sigma_{m,i}) \times p(s_{ijk}|\mu_{m,i}, \sigma_{m,i}) \times p(m_{ijk}|\gamma) \times p(f_{ij}|\mu_{m,i}, \sigma_{m,i}) \times p(y_i) \times p(b_l). \tag{23}$$

In particular

$$p(z_{ijkl}|y_{ijkl}, \sigma_{\varepsilon,i}) = \begin{cases} 0 & \text{with probability } 1 - p_{ijkl} \\ \text{Normal}(y_{ijkl}, \sigma_{e,i}) & \text{with probability } p_{ijkl}. \end{cases} \tag{24}$$

Posterior distribution for all model parameters were estimated using the variational inference [2] method implemented in the Python package 'pymc3' [4].
